# *In vitro* differentiation of mouse pluripotent stem cells into glucocorticoid-producing adrenocortical cells

**DOI:** 10.1101/2023.12.22.572808

**Authors:** Ioannis Oikonomakos, Melina Tedesco, Fariba Jian Motamedi, Mirko Peitzsch, Serge Nef, Stefan Bornstein, Andreas Schedl, Charlotte Steenblock, Yasmine Neirijnck

## Abstract

Directed differentiation of pluripotent stem cells into specialized cell types represents an invaluable tool for a wide range of applications. Here, we have exploited single-cell transcriptomic data to develop a step-wise *in vitro* differentiation system from mouse embryonic stem cells into adrenocortical cells. We show that during development the adrenal primordium is embedded in an extracellular matrix containing tenascin and fibronectin. Culturing cells on fibronectin during differentiation increased the expression of the steroidogenic marker NR5A1. Furthermore, 3D cultures in the presence of PKA-pathway activators led to the formation of aggregates composed of different cell types expressing adrenal progenitor or steroidogenic markers, including the adrenocortical specific enzyme *Cyp21a1*. Importantly, *in vitro* differentiated cells secreted the *zona Fasciculata* specific corticosterone in response to cAMP activators, but not gonadal hormones, thus confirming the specificity of differentiation towards the adrenal lineage.

**Highlights:**

- Mouse adrenal progenitors differentiate in a tenascin/fibronectin-rich extracellular matrix
- Stepwise differentiation of ES cells yields NR5A1+ adrenocortical cells
- 3D aggregates produce glucocorticoids in response to cAMP signalling

## Introduction

The adrenal cortex is a vital component of the body’s regulatory system regulating blood pressure, metabolism and the response to stress and infections through the release of steroid hormones. Steroids are produced in highly specialized zones that are arranged in concentric rings including the mineralocorticoid-producing zona glomerulosa (zG), the glucocorticoid-producing zona fasciculata (zF), and the androgen-producing zona reticularis (zR). Mouse adrenals lack a functional zR, but have a juxtamedullary X-zone, a remnant of the fetal cortex that disappears in males at puberty and in females after the first pregnancy ^1^. Given the important function of adrenals, it is not surprising that adrenal insufficiencies, such as Addison’s disease and congenital adrenal hyperplasia (CAH) ^2–4^, can be life-threatening if left untreated. Hormone therapy is the primary treatment but maintaining adequate hormone levels can be challenging, leading to hormonal fluctuations that can affect the quality of life and increase the risk of adrenal crisis ^5,6^.

A potential cure for adrenal insufficiency may be the transplantation of functional adrenocortical cells capable of responding to adrenocorticotropic hormone (ACTH) to ensure an adequate release of glucocorticoids. Functional steroidogenic cells have previously been generated by forced expression of the transcription factor NR5A1 ^7–9^, a master regulator of steroidogenesis ^10,11^. However, overexpression of NR5A1 results in abnormal steroid hormone production, thus limiting the use of such a system for therapeutic approaches. The development of protocols that allow directed differentiation of pluripotent stem cells into adrenocortical cells could potentially overcome this problem. In a recent study, human induced pluripotent stem cells (hIPSCs) were used to generate steroidogenic cells, but instead of glucocorticoids, these cells produced dehydroepiandrosterone (DHEA), a precursor of androgens and oestrogens ^12^. *In vitro* differentiation of pluripotent stem cells into glucocorticoid-producing cells has not yet been reported.

To develop an *in vitro* differentiation protocol, a detailed knowledge of adrenal development is essential. Adrenocortical progenitors are mesoderm-derived and emerge from the adrenogonadal primordium (AGP), a structure traditionally defined by *Nr5a1* expression and located at the interface of the intermediate mesoderm (IM) and the lateral plate mesoderm (LPM) ^11^. Recent advances using single-cell transcriptomic analysis in mice ^13,14^ and humans^15^ have permitted further insights into the adrenal origin. In particular, these studies have revealed that adrenocortical cell fate is specified prior to *Nr5a1* expression, at the anterior end of the developing urogenital ridge.

Here, we have leveraged these findings to develop a stepwise differentiation protocol of mouse pluripotent embryonic stem cells (mESCs) into adrenocortical cells *via* anterior intermediate/lateral plate mesodermal progenitors. Importantly, when cultured in 3D, the aggregates endogenously express key markers of adrenocortical cells and secrete the steroids corticosterone and 11-dehydrocorticosterone.

## Results

### Directed differentiation of mESCs into anterior intermediate/lateral mesoderm induces expression of early adrenocortical progenitor markers

Early mammalian development involves the differentiation of ESCs into epiblast stem cells (EpiSCs), transitioning from naïve to primed pluripotency. Epiblastic cells further undergo an epithelial to mesenchymal transition to give rise to the primitive streak (PS), a transient embryonic structure comprising mesendodermal precursors that can be induced by canonical Wnt signalling both *in vivo* and *in vitro* ^16^ (**Fig.1A**). To convert mESCs into EpiSCs, we first treated mESCs with activin and FGF2 for 2 days, as previously described ^17^, resulting in upregulation of the EpiSC marker *Fgf5* (**Fig. S1A, B**). We next induced Wnt/β-catenin signalling with the GSK3-α/β inhibitor Chiron (CHIR99021) for 2 days ^18,19^ and observed induction of the PS as evidenced by robust upregulation of *Brachyury* (*T*) at day 4 of differentiation (**Fig.S1B**).

**Figure 1.**
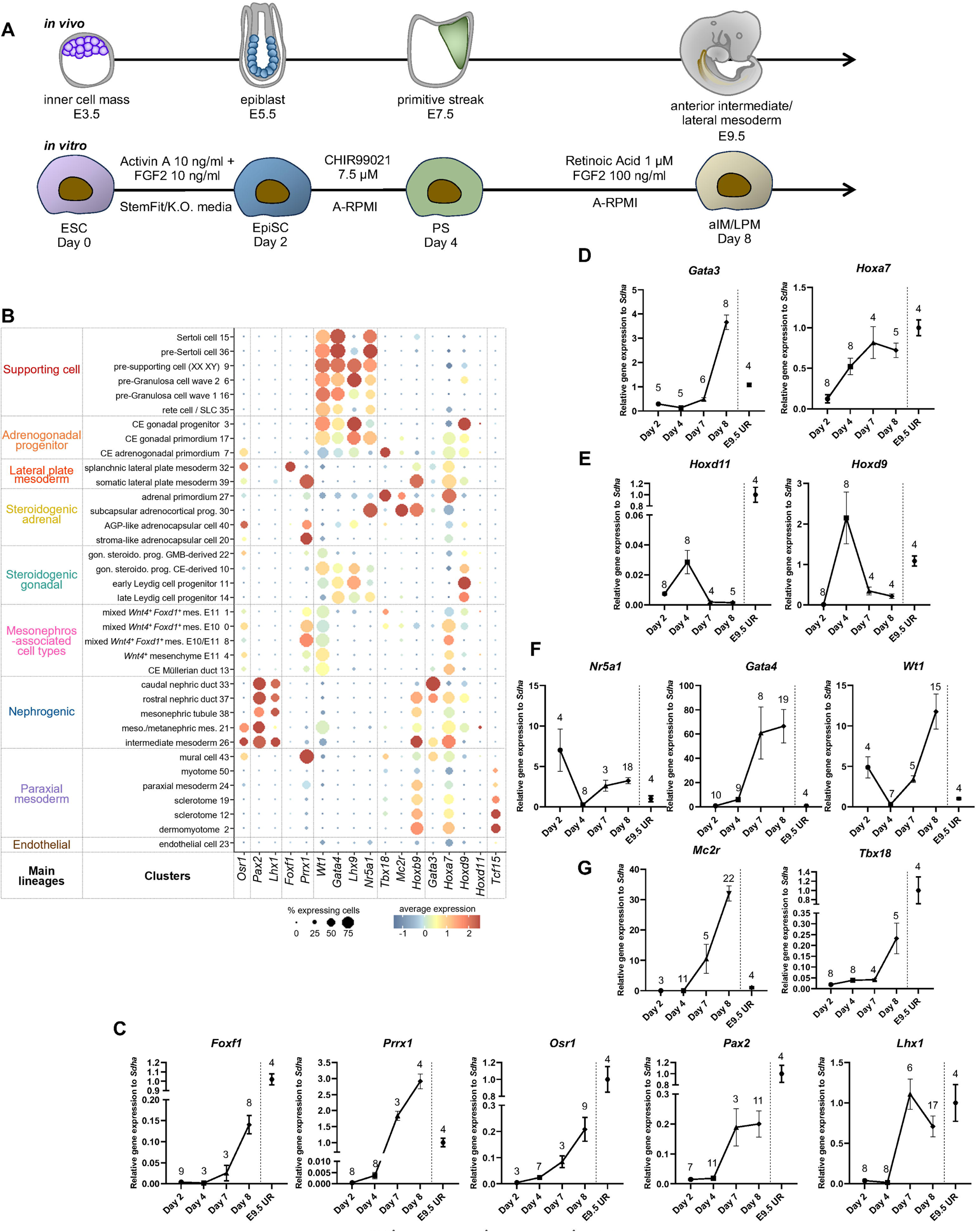
Differentiation of mESCs to anterior intermediate/lateral mesoderm is associated with early adrenocortical marker expression. (A) Schematic overview of adrenal cortex development *in vivo* (top panel) and stepwise mESC differentiation *in vitro* (bottom panel). Cells were treated two days with Activin A and FGF2, two days with CHIR99021 and four days with retinoic acid and FGF2 (see also experimental procedures). aIM/LPM, anterior intermediate/lateral plate mesoderm; E, embryonic day; EpiSC, epiblast stem cell, ESC, pluripotent embryonic stem cell; PS, primitive streak. (B) Dot plot showing expression of selected marker gene (x axis) per cluster (y axis) of the mesodermal-derived cell lineages from Neirijnck et al.^13^. The size of the dot represents the percentage of cells in the cluster expressing the gene, and the colour indicates the level of expression (log normalized counts). Cluster annotation and major cell lineages are displayed on the left. AGP, adrenogonadal primordium; CE, coelomic epithelium; E, embryonic day; GMB, gonad/mesonephros border; gon. steroido. prog., gonadal steroidogenic progenitor; mes., mesenchyme; meson., mesonephric; SLC, supporting-like cell. (C-G) RT-qPCR for marker genes of intermediate and lateral plate mesoderm (*Foxf1*-*Pax2* **; *Prrx1-Osr1*\*\*\*; *Lhx1*\*\*\*\*) (C), antero-posterior patterning of the urogenital ridge (*Hoxd9-Hoxd11-Hoxa2*\*\*; *Gata3*\*\*\*\*) (D and E), early adrenogonadal progenitors (*Gata4*\*\*; *Wt1*\*\*\*; *Nr5a1*\*\*\*\*) (F) and early adrenocortical progenitors (*Tbx18*\*\*; *Mc2r*\*\*\*\*) (G), during the progress of *in vitro* differentiation. Data depict relative mRNA expression levels, presented as mean ±SEM values of independent replicates (number indicated on the graphs) and normalized on E9.5 mouse urogenital region (E9.5 UR, pools of 4 to 6 embryos). Welch’s ANOVA test was performed resulting in significant difference among means. **=p<0.01; ***=p<0.001; ****=p<0.0001. See also Fig S1.

Previous work has shown that hIPSCs-derived PS cells spontaneously differentiate into definitive endoderm and mesodermal subtypes, a cell fate that can be redirected towards IM by the exogenous addition of fibroblast growth factor 2 (FGF2) ^19^, and anteriorized by the addition of retinoic acid (RA) ^18^. We tested whether treatment of mESCs-derived PS cells with these factors would direct specification towards the anterior IM and/or LPM, which would provide adequate conditions for the specification into adrenocortical progenitors. To allow fine-tuning of differentiation protocols, we first analysed our recently published single-cell transcriptomic atlas ^13^ to establish a set of suitable developmental markers that highlight specific lineages and stages (**Fig.1B**). Consistent with earlier reports ^20^, *Osr1* was found to be expressed at early stages in IM (cluster (C) 26), nephrogenic mesenchyme (C21), mesonephros-associated cell types (C1, C0, C8, C4 and C13) and LPM (C32 and C39), but was absent in paraxial mesoderm (C2, C12, C19 and C24). We therefore considered this gene as a marker for both IM and LPM. In contrast, expression of *Pax2*/*Lhx1* ^21,22^ and *Foxf1/Prrx1* ^23,24^ are mutually exclusive in the IM and LPM lineages, respectively, confirming previous reports. Early adrenogonadal progenitors (C7) expressed the transcription factors *Wt1*, *Gata4, Lhx9, Tbx18* and low levels of *Nr5a1.* As the cells progressed through the adrenal (C27 and C30) or gonadal (C17 and C3) lineages, they exhibited distinct transcriptional profiles with expression of *Tbx18/Mc2r/Hoxb9* and *Wt1/Gata4/Lhx9*, respectively, while *Nr5a1* was upregulated in both lineages. In addition, the expression of the anterior *Hoxa7* and posterior *Hoxd9* genes distinguish the adrenal primordium (C27) from the gonadal primordium (C17), respectively, reflecting lineage specification along the anteroposterior axis ^13^. Treatment of mESC-derived PS cells with FGF2 and RA for 4 days (day 8 of the differentiation protocol) resulted in strong expression of *Osr1*, *Pax2, Lhx1, Foxf1* and *Prrx1* (**Fig.1C**), indicating differentiation into IM and LPM cells. Paraxial mesoderm (*Tcf15)* and endoderm (*Pdx1*) markers were not upregulated (**Fig.S1C**) thus confirming the specificity of the differentiation. Further analysis revealed higher expression levels of anterior (*Gata3/Hoxa7)* than posterior (*Hoxd9/Hoxd11*) markers (**Fig.1D-E**)^18^ suggesting that the cells were adopting an anterior intermediate/lateral plate mesoderm (aIM/LPM) fate. Interestingly, this was concomitant with the induction of the adrenogonadal progenitor markers *Wt1, Gata4* and *Nr5a1* (**Fig.1F**) and an increase in the adrenocortical progenitor marker genes *Tbx18, Hoxb9*, and *Mc2r*, encoding the ACTH receptor (**Fig.1G** and **Fig.S1D**). By contrast the gonadal-enriched gene *Lhx9* was not found to be induced suggesting differentiation towards the adrenal rather the gonadal lineage (**Fig.S1D**). The steroidogenic marker *Nr5a1,* which is expressed at low levels in ESCs ^25^, but is lost during differentiation to the PS state, showed only a mild induction upon FGF2 and RA treatment, in contrast to *Wt1* whose expression kept increasing over time (**Fig.S1E**).

### Extracellular matrix promotes endogenous NR5A1 expression

We ^13^ and others ^26^ have previously shown that early adrenocortical cells are specified prior to *Nr5a1* expression. The expression of the adrenocortical-enriched genes *Tbx18, Hoxa7, Hoxb9* and *Mc2r* at day 8 of differentiation, but absence of high levels of *Nr5a1,* suggested that the cells had committed towards the adrenocortical lineage but had failed to fully differentiate. To drive the aIM/LPM cells towards the steroidogenic fate, we aimed to mimic the *in vivo* adrenocortical environment with a focus on extracellular matrix (ECM) proteins. Analysis of our single-cell atlas revealed that adrenogonadal progenitors specifically express *tenascin C* (*Tnc)*, while several cell types expressed *fibronectin 1* (*Fn1)* (**Fig.2A**). Immunostaining confirmed TNC expression in NR5A1^+^ cells at embryonic day (E) 10.5 (**Fig.2B**), whereas FN1 was more widely expressed (**Fig.2C**). Interestingly, *Tnc* expression was strongly induced (∼100 fold) from day 4 to day 8 in our *in vitro* system, further supporting the correct fate of the cells towards the adrenogonadal lineage. *Fn1* expression remained stable during this period (**Fig.2D**).

**Figure 2.**
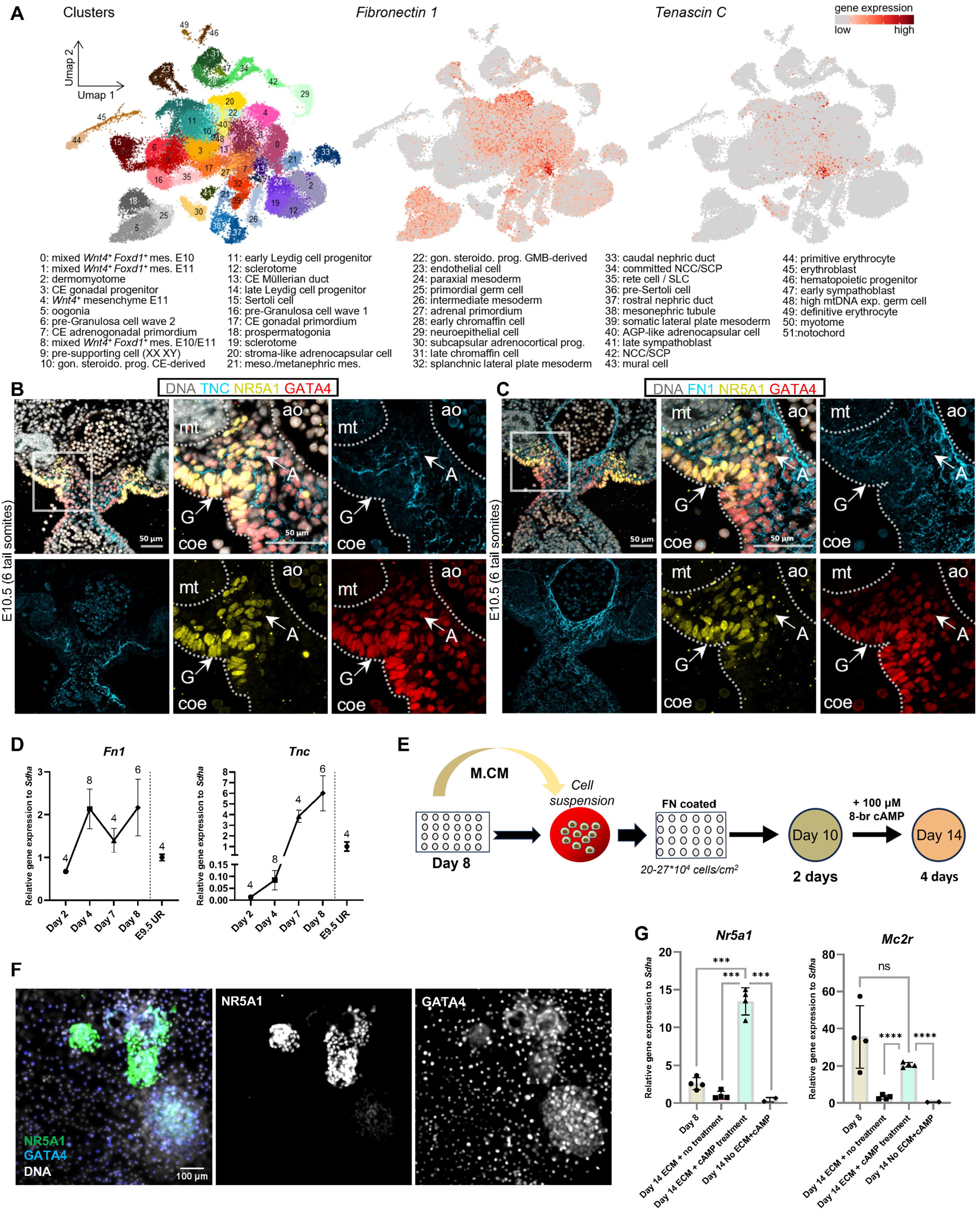
Adrenocortical progenitors differentiate in a fibronectin/tenascin C rich environment. (A) UMAP visualization of the 72,273 transcriptomes from ^13^ coloured by clusters and by expression levels of *fibronectin 1* (*Fn1*) and *tenascin C* (*Tnc*). Cluster annotation is shown on the bottom. AGP, adrenogonadal primordium; CE, coelomic epithelium; GMB, gonad/mesonephros border; gon. steroido. prog., gonadal steroidogenic progenitor; mes., mesenchyme; meson., mesonephric; mtDNA exp., mitochondrial DNA expressing; NCC, neural crest cell; SLC, supporting-like cell; SCP, Schwann cell precursor. (B) Co-labeling for GATA4, NR5A1, and TNC (immunostaining) in E10.5 embryos (transverse sections), revealing TNC expression in mesenchymal cells underlying the coelomic epithelium, including the adrenal primordium. Nuclei are stained with Hoechst. A, adrenal field, ao, aorta; coe, coelomic cavity; G, gonadal field; mt, mesonephric tubule. Scale bars, 50 μm. (C) Co-labeling for GATA4, NR5A1 and FN1 (immunostaining) in E10.5 embryos (transverse sections), revealing broad FN1 expression in mesenchymal cells. Nuclei are stained with Hoechst. A, adrenal field, ao, aorta; coe, coelomic cavity; G, gonadal field; mt, mesonephric tubule. Scale bars, 50 μm. (D) RT-qPCR for *Fn1* (*) and *Tnc (**)* during the progress of *in vitro* differentiation. Data depict relative mRNA expression, presented as mean ±SEM values of independent replicates (number indicated on the graphs) and normalized on E9.5 mouse urogenital region (E9.5 UR, pools of 4 to 6 embryos). Welch’s ANOVA test was performed resulting in significant difference among means. *=p<0.05; **=p<0.01). (E) Schematic overview of aIM/LPM cells (day 8) differentiation on FN-coated plates. Cells were treated 2 days with M.CM and 2 days with 8-br-cAMP (see also experimental procedures). (F). Co-labeling for NR5A1 and GATA4 in Day 14 differentiated cells, revealing NR5A1^+^ colonies. Nuclei are stained with Hoechst. Scale bars, 100 μm. (G) RT-qPCR for *Nr5a1* and *Mc2r* on aIM/LPM cells (day 8) and on day 14 cells grown on FN1 (ECM treatment) and/or 8-br-cAMP supplementation (cAMP treatment) (see also experimental procedures). Data depict relative mRNA expression, presented as mean ±SD values of independent replicates (n=2 to 4) and normalized on E9.5 mouse urogenital region (pools of 4 to 6 embryos, n=4). Statistical analysis was performed using two-tailed unpaired Welch’s t-test. ns., not significant; **=p<0.01; ***=p<0.001; ****=p< 0.0001.

We next aimed to replicate the FN1-rich environment in our *in vitro* system. Day 8 cells were detached from their original collagen- and laminin-based ECM (Geltrex) and replated onto plates coated with FN1. To wean the cells from their familiar environment, day 8 conditioned media (M.CM) was added during the introduction of the new ECM ^27,28^. *Nr5a1* expression has been shown to be stimulated by PKA signalling ^29,30^. We therefore added 8-Br-cAMP (a stable analogue of cAMP) after two days in the new culture conditions (**Fig.2E**). After a further four days of differentiation, the colonies contained cells with strong NR5A1 expression surrounded by GATA4^+^ cells (**Fig.2F**). RT-qPCR analysis confirmed that the combination of FN1 coating and cAMP supplementation resulted in a strong increase in *Nr5a1* expression compared to FN1 coating or cAMP supplementation alone (**Fig.2G**). Expression of *Mc2r* was maintained under FN1/cAMP conditions but decreased in the presence of FN1 or cAMP alone.

### 3D cultures of *in vitro*-derived aIM/LPM cells induce adrenocortical-specific steroidogenic gene expression

In other systems, 3D cultures have been shown to be more conducive to differentiation by providing molecular cues from neighbouring cells as well as mechanical stimulation ^31,32^. We therefore tested the aggregation of aIM/LPM cells in ultra-low attachment plates, or in microwell plates, the latter allowing the formation of hundreds of small spheroids in a single well. After two days in M.CM (day 10 of differentiation), aggregates were treated with or without 8-Br-cAMP for 5 days (day 15 of differentiation) (**Fig.3A**). Quantitative PCR analysis at days 8, 10, 12 and 15 was performed to evaluate the expression dynamics of adrenocortical specific markers. Culturing in microwell plates generally resulted in higher activation of steroidogenic genes than in ultra-low attachment plates **(Fig.S2A, B)** and this method was chosen for all further experiments.

**Figure 3.**
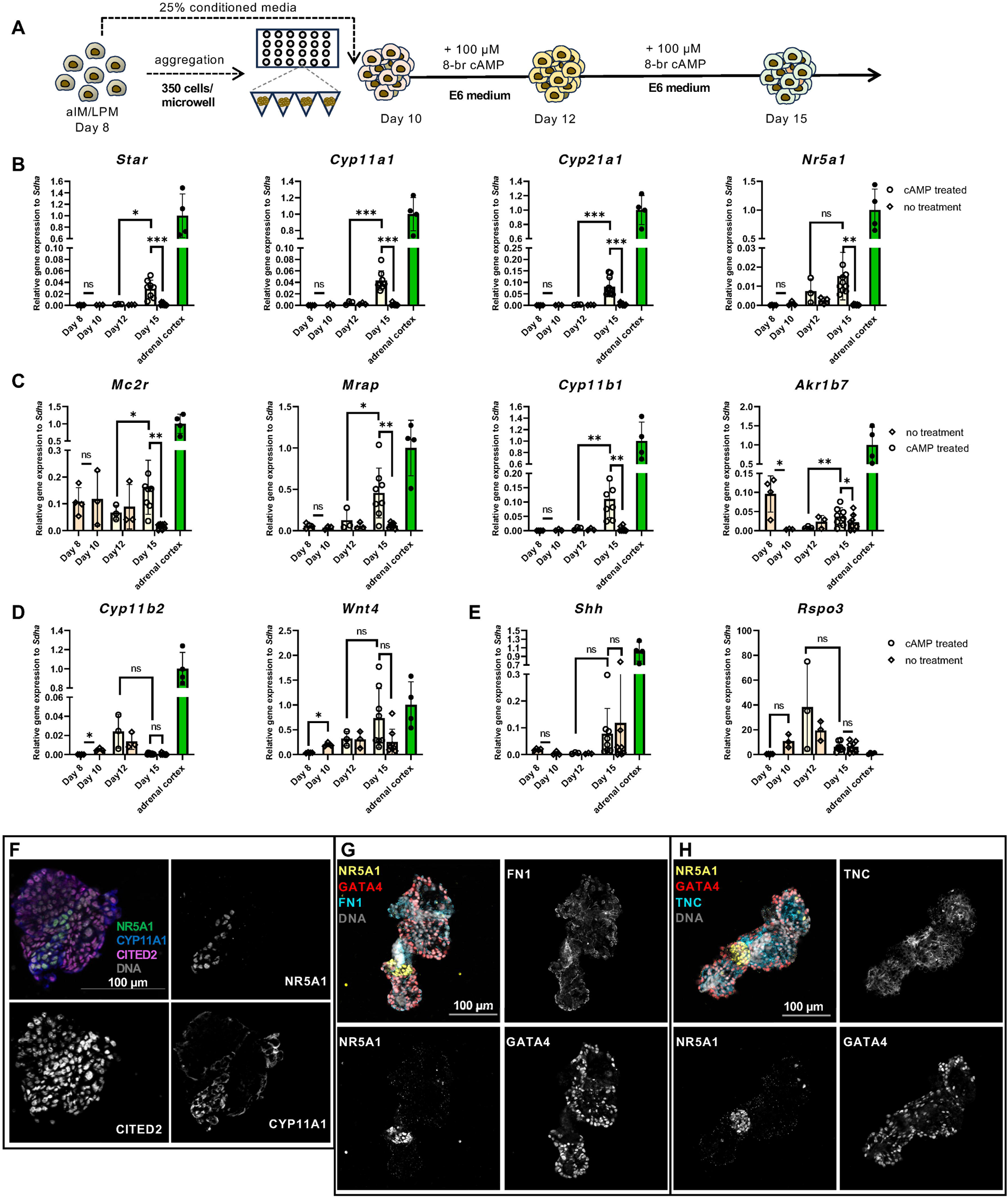
Differentiation of aIM/LPM cells with 3D assembly leads to a steroidogenic fate. (A) Schematic overview of aIM/LPM cells differentiation. Cells are aggregated on microwell plates and treated for 2 days with M.CM and 5 days with 8-br-cAMP (see also experimental procedures). (B-E) RT-qPCR analysis for the steroidogenic genes *Star*, *Cyp11a1*, *Cyp21a1*, *Nr5a1* (B), the corticosterone production-related genes *Mc2r*, *Mrap*, *Cyp11b1* and the zona Fasciculata marker *Akr1b7* (C), the aldosterone production-related genes *Cyp11b2* and *Wnt4* (D), and the adrenocortical progenitor markers *Rspo3* and *Shh* (E), during the progress of *in vitro* differentiation. Data depict relative mRNA expression, presented as mean ±SD values of independent replicates (Day8 n=4, Day10 n=3, Day12 treated and not treated n=3, Day15 treated and not treated n= 8), and normalized on adult adrenal cortex+capsule (n=4). Statistical analysis was performed using two-tailed unpaired Welch’s t-test. ns., not significant; *=p< 0.05; **=p<0.01; ***=p<0.001. (F-H) Co-labeling for NR5A1, CYP11A1, CITED2 (F), NR5A1, GATA4, FN1 (G) and NR5A1, GATA4, TNC (H) on Day 15 paraffin-embedded aggregates. Nuclei are stained with Dapi or Hoechst. Scale bars, 100 μm.

cAMP stimulation led to a significant increase in *Nr5a1,* the cholesterol transporter *Star*, the steroidogenic enzyme *Cyp11a1* and the adrenal specific enzyme *Cyp21a1,* although the levels remained lower than in adrenal cortex microdissected from adult mice (**Fig.3B**). Medium without 8-Br-cAMP supplementation resulted in significantly lower expression of these genes. Analysis of markers specific for glucocorticoid-producing zF cells revealed cAMP-dependent induction of *Cyp11b1* and *Mc2r-accessory protein* (*Mrap*), and cAMP-dependent maintenance of *Mc2r* expression (**Fig.3C**), whereas *Akr1b7* was expressed at day 15 independently of the treatment (**Fig.3C**). Markers of mineralocorticoid-producing zG cells, *Cyp11b2*, *Wnt4* and *Dab2* were also induced during differentiation, but this was independent of cAMP treatment (**Fig.3D** and **Fig.S3A**). Moreover, the expression of the key gene for mineralocorticoid production (*Cyp11b2*) was far lower than *in vivo.* Interestingly, *Rspo3* and *Shh*, which are expressed in capsular or subcapsular progenitors of the definitive adult cortex ^33,34^, remained highly expressed with and without cAMP stimulation (**Fig.3E**). This suggests that both differentiated steroidogenic and progenitor cell populations persist under these culture conditions. Importantly, no significant levels of the gonadal progenitor marker *Lhx9*, the supporting cell markers *Wt1, Sox9* and *Foxl2* ^35^, neither of testicular steroidogenic enzyme *Hsd17b3* ^36^ were observed (**Fig. S3B-D**), indicating specificity of differentiation towards the adrenal lineage.

To gain further insight into the architecture of the aggregates, we performed immunostaining for NR5A1, CYP11A1 and the adrenal primordium marker CITED2 ^37^ (**Fig.3F**). NR5A1^+^/CYP11A1^+^ cells were found preferentially embedded in NR5A1^-^/CITED2^+^ cells, indicating that both differentiated and non-steroidogenic progenitor cells are present in the aggregates, in agreement with our qPCR analysis. Moreover, immunostaining for FN1 and TNC showed an enrichment of these ECM proteins in the aggregates (**Fig.3G, H**), indicating that 3D cultured cells produce ECM proteins that are also found *in vivo*.

### Aggregates produce glucocorticoids

The expression of genes involved in the processing of steroidogenic enzymes and the presence of *Cyp11b1* suggested that the *in vitro*-differentiated aggregates were able to produce zF-specific steroids (**Fig.4A**). To address this possibility, we collected medium on day 15 and performed steroid hormone analysis by liquid chromatography with tandem mass spectrometry (LC-MS/MS) ^38^. The 8-Br-cAMP treated samples showed the presence of the main mouse glucocorticoids corticosterone (∼8 ng/ml), 11-dehydrocorticosterone (11-DHC) (∼12 ng/ml) and 11-deoxycorticosterone (DOC) (∼2 ng/ml) in the media (**Fig.4B**). No mineralocorticoids (aldosterone) were detected (**Table S1**). Importantly, sex steroids such as testosterone and dihydrotestosterone, were barely detectable (**Fig.4B**), indicating a high specificity of adrenocortical steroid production in the *in vitro*-differentiated aggregates. Furthermore, ELISA analysis indicated that the initiation of corticosterone production occurred after day 12 of differentiation (**Fig.4C**), a finding that is consistent with the late induction of steroidogenic genes (**Fig.3B-C**).

**Figure 4.**
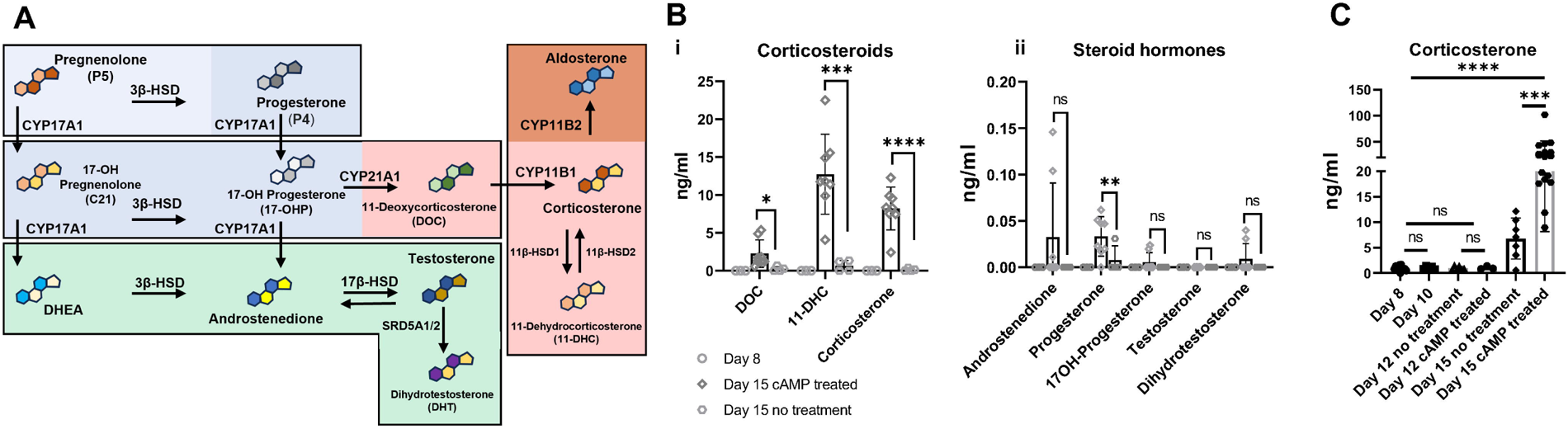
*In vitro*-differentiated aggregates produce glucocorticoids. (A) Illustration of the steroid production pathway in mice, delineating the key genes responsible for synthesizing progestogens (blue box), corticosteroids (red box) and androgens (green box). (B) LC-MS/MS analysis for corticosteroids (i) and progestogens/androgens (ii) on M.CM (Day 8) and aggregates medium (Day 15) treated or not with 8-br-cAMP. Data depict steroid concentrations as ng/ml of medium, presented as mean ±SD values of independent replicates (Day8 n= 3, Day15 cAMP treated n= 8, Day15 not treated n= 4). Statistical analysis was performed using two-tailed unpaired Welch’s t-test. ns.=not significant; *=p<0.05; ***=p<0.001, ****=p<0.0001. DOC, 11-Deoxycorticosterone; 11-DHC, 11-Dehydrocorticosterone. See also Table S1. (C) ELISA measurement of corticosterone levels in M.CM (Day 8) and medium collected from aggregate cultures (Days 10, 12 and 15) treated or not with 8-br-cAMP. Data depict corticosterone concentrations as ng/mL of medium, presented as means ±SD values of independent experiments (Day8 n= 11, Day10 n= 5, Day12 cAMP treated n= 3, Day12 not treated n= 6, Day15 cAMP treated n= 17, Day15 not treated n= 7). Statistical analysis was performed using two-tailed unpaired Welch’s t-test. ns.=not significant; *=p < 0.05; ***=p < 0.001; ****=p < 0.0001.

## Discussion

Cellular model systems are essential for studying the molecular and cellular functions of any given cell type. Currently, *in vitro* systems for studying adrenocortical cells are limited to primary cells, which can only be maintained in culture for a short time, and a handful of cell lines mainly derived from cancer patients. The protocol developed here allows the differentiation of mouse ES cells into adrenocortical-like cells that produce steroidogenic enzymes and secrete the main mouse glucocorticoids corticosterone and 11-deoxycorticosterone. The relatively simple protocol appears to be robust as the differentiation has been achieved using different ES cell strains (C57BL/6, R129) in two different institutes, although some adjustments to the concentrations of the GSK3-α/β inhibitor Chiron are required to optimize mesendoderm production and cell survival for different cell lines.

Our protocol follows the major steps of embryonic differentiation and employs growth factors and signalling pathways that have been used previously for mesoderm induction. However, cellular differentiation is not only driven by diffusible signalling molecules ^39–42^ but is also influenced by the ECM ^43,44^. By analysing the data from of our previously published single-cell atlas ^13^, we discovered the presence of several ECM proteins expressed during adrenal specification, including TNC and FN1. The expression of FN1 in the developing adrenal cortex has been reported previously ^45–47^. The immunostaining in mouse embryos presented here confirmed that TNC expression is highly specific to the adrenal, but not gonadal lineage, at this developmental stage. Previous studies have shown that FN1 and TNC interact with each other ^48–50^ to provide an invasive environment for cancer cells ^51^, and it is tempting to speculate that the interaction between these molecules may play a similar role in supporting differentiation and the migration of adrenal progenitors from the coelomic epithelium to a more medial position. Interestingly, culturing cells in 2D on FN1 increased NR5A1 expression. TNC exists in many different isoforms that appear to have different functions ^52,53^. It will be interesting to conduct further studies to determine which variants are produced and how they influence the development and maturation of the adrenal cortex. Of note, 3D aggregates express TNC and FN1 suggesting that our culture conditions recapitulate the *in vivo* environment.

*In vitro* differentiation of pluripotent stem cells rarely produces pure cell populations, but rather results in diverse cell types that often reflect various degrees of differentiation. We observed a similar diversity of cells in our 3D culture system with differentiated cells characterized by high levels of NR5A1 and CYP11A1 embedded in cells expressing GATA4, a marker of the AGP ^54^, and CITED2, a progenitor marker of the adrenal gland ^37^. Further optimization of culture conditions may allow a full conversion of these progenitors into steroid producing cells.

Adrenals and gonads have long been thought to derive from a common primordium characterized by the expression of NR5A1 ^55^. Recent single-cell sequencing studies from our lab and others have challenged this model, and suggest that adrenals and gonads are specified independently, with the adrenal primordium arising in a more anterior position ^13,15^. The expression of the anterior markers *Gata3* and *Hoxa7* and the lack of strong induction of the posterior markers *Hoxd9* and *Hoxd11* at day 8 of differentiation suggest that our protocol does indeed targets the anterior part of the urogenital ridge. The absence of gonadal markers (*Foxl2*, *Sox9, Hsd17b3*) and gonadal hormones and the presence of *Cyp21a1* confirms the specificity of adrenocortical differentiation.

Mineralocorticoids and glucocorticoids are produced by distinct zones of the mouse adrenal cortex. In our system we only detected corticosterone, 11-dehydrocorticosterone and 11-deoxycorticosterone suggesting that cells have adopted a zF-like differentiation status. Glucocorticoid production is tightly controlled by the HPA axis, which induces PKA signalling in zF cells and previous research has shown that zF differentiation is also dependent on cAMP activation ^29^. It is therefore not surprising that the addition of 8-Br-cAMP in the final steps of our protocol results in cells that produce glucocorticoids rather than mineralocorticoids. When analyzing the expression levels of the zG specific marker *Cyp11b2* we noticed an initial increase at days 10-12 during differentiation, which may suggest that progenitor cells at this stage are poised to differentiate into either zG-or zF-like cells. It will be interesting to test whether altering culture conditions, perhaps by reducing cAMP levels and adding β-catenin activators such as R-spondins ^33^ and WNT4 ^56^, can modify the differentiation path towards the zG and result in mineralocorticoid-producing cells.

In summary, we report here the generation of functional adrenocortical steroidogenic cells *in vitro*. We envisage that the system presented here will become a powerful tool to study normal and abnormal development and differentiation *in vitro.* Furthermore, the impact of mutations on steroidogenesis and other cellular functions can be tested. As the protocol is relatively simple it may be suitable for efficient drug testing. Future research will focus on adapting the protocol to human iPSCs, which in the long term may allow the development of cell replacement therapies for patients suffering from adrenal insufficiency or congenital adrenal hyperplasia.

## Experimental procedures

### Resource availability

#### Corresponding authors

Further information and requests for resources and reagents should be directed to and will be fulfilled by the corresponding authors.

#### Materials availability

All unique/stable reagents generated in this study are available from the lead contact.

#### Data and code availability

Single-cell RNA-seq data have been previously published ^13^ and are available at the NCBI Gene Expression Omnibus database (GEO: GSE156176).

### Single-cell transcriptomic analysis

Single-cell gene expression data have been computed from the previously published atlas of mouse adrenogonadal development ^13^ using Seurat2.3.4 ^57^ on the entire dataset (72,273 cells) or only on mesodermal derivatives (59,619 cells) as indicated in the legend of the figures.

### Animals

All animal work was conducted according to ethical guidelines of the Direction Générale de la Santé of the Canton de Genève (experimentation ID GE/57/18) and the ethical and research board of the Landesdirektion Sachsen (experimentation ID TVvG 10/2022). For paraffin embedding of embryos, the *Nr5a1-eGFP^Tg^*mouse strain ^58^ maintained on a CD1 genetic background was used. E10.5 (8 ± 2 tail somites (ts)) embryos from timed matings (day of vaginal plug = 0.5) were collected and processed for paraffin embedding. For RNA preparation, the C57BL/6J-6N strain was used. Urogenital ridges from E9.5 embryos (22 ± 2 somites), adrenal glands from E14.5 embryos and adrenal cortex+capsule ^59^ from 12-14 weeks old mice were collected and snap frozen until use.

### mESCs maintenance

R1 mESCs ^60^ (ATCC SCRC-1011) were used in this study. mESCs were cultured in feeder-free conditions on tissue-culture treated 6-well plates (Falcon® 353046) coated with 0.1% gelatine (Sigma G1393), in KnockOut DMEM (Gibco™ 10829018) supplemented with 15% KnockOut serum replacement (Gibco™ 10828028), Glutamax (Gibco™ 35050061), non-essential amino acids (Gibco™ 11140035), 50 µM 2-Mercaptoethanol (Gibco™ 31350010), 1000 U/ml ESGRO recombinant mouse LIF protein (Sigma ESG1106), 2 μM PD0325901 (Selleckchem S1036) and 3 μM CHIR9902 (Sigma SML1046) ^61^, at 37°C with 5% CO_2_. Cells were passaged using 0.25% Trypsin-EDTA solution (Gibco™ 25200056) and Newborn Calf Serum (Gibco™ 16010159).

### Differentiation of mESCs to aIM/LPM cells

Before differentiation, cells were kept in culture and passaged at least once. mESCs were dissociated as mentioned before and seeded at 1.8*10^4^ cells/cm^2^ density on new 24-well tissue-culture plates (Falcon® 353047) coated with 1% Geltrex (Gibco™ A1413302) in DMEM-F12 (Gibco™ 11320033) at 37°C for 3-4h. For EpiSCs differentiation, cells were cultured from day 0 to day 2 in 1:1 ReproFF2 (REPROCELL RCHEMD006) or StemFit™ Basic04CT (REPROCELL ASB04CT): maintenance media (see above) without LIF, PD0325901 and CHIR9902, and supplemented with 10 ng/ml recombinant Activin A (STEMCELL^TM^ 78001) and 10 ng/ml recombinant FGF2 (Peprotech 100-18B). ReproFF2 or StemFit media resulted in comparable gene induction (data on request). For PS differentiation, cells were cultured from day 2 to day 4 in advanced RPMI 1640 (A-RPMI) (Gibco™ 12633012) supplemented with GlutaMax and 7.5 μM CHIR99021. For aIM/LPM differentiation, cells were washed with A-RPMI and cultured from day 4 to day 8 in A-RPMI supplemented with 1 μM retinoic acid (Sigma R2625) and 100 ng/ml FGF2. All differentiation procedures were performed at 37°C with 5% CO2. Media were refreshed every day keeping ∼10% of initial media in the well.

### Plating and differentiation on fibronectin

Wells of a 24 flat and clear bottom plate (Ibidi 82426) were coated with 1 μg/ml FN1 (Corning™ 354008) in DPBS, centrifuged at ∼600 g for 2 min, incubated overnight at 4°C and brought to 37°C for an hour before plating the cells. The mesodermal conditioned medium (M.CM) from day 8 was first gathered and filtrated. Cells were dissociated using 1:1 Trypsin:DPBS and FBS (Gibco™ 10082147) and seeded at a 20-27*10^4^ cells/cm^2^ density on the FN1-coated wells. Cells were cultured from day 8 to day 10 in A-RPMI supplemented with 25% M.CM, and from day 10 to day 15 in A-RPMI supplemented with 100 μM 8-br-cAMP (Tocris 1140). Media were refreshed at day 12 keeping ∼20% of the initial medium in the well.

### Aggregate formation and differentiation

Aggregates were formed in either Ultra Low attachment (ULA) plates (Corning® Costar® CLS7007) or in microwells (SphericalPlate 5D® (Kugelmeiers SP5D-24W) or AggreWell™400 (STEMCELLTM 34411)). M.CM was harvested and cells were dissociated as described above. 2*10^4^, 3*10^5^ and 4.2*10^5^ cells were seeded per well of ULA, SphericalPlate 5D and AggreWell™ 400 plates, respectively. For ULA plates, aggregation was achieved by centrifugation for 2 min at ∼145 g. In microwell plates, aggregates formed spontaneously. Aggregates were cultured from day 8 to day 10 in A-RPMI or TeSR™-E6 (StemCell Technologies 05946) medium supplemented with 25% M.CM. At day 10 of differentiation, an equal volume of media (A-RPMI or TeSR™-E6) supplemented with 200 μM 8-br-cAMP was added, to reach a final concentration of 100 µM 8-br-cAMP. At day 12 of differentiation, half of the medium was refreshed with new media supplemented with 100 µM 8-br-cAMP.

### RNA isolation and RT-qPCR

Total RNA was extracted using the RNeasy MicroKit or the RNeasy MiniKit (Qiagen 74004 and 74106) according to the manufacturer’s protocol. RNA was reverse transcribed using SuperScript™ III Reverse Transcriptase (Invitrogen 18080093) according to the manufacturer’s protocol, in combination with random primers (Invitrogen 48190011), dNTPs (Invitrogen 18427013) and RNaseOUT™ (Invitrogen 10777019).

Quantitative PCR was performed with the LightCycler 480 SYBR Green I Master (Roche 04707516001) on a Real-Time PCR System (LightCycler® 480 (Roche) or CFX Connect (BIO-RAD)), with 40 amplification cycles and 60°C annealing temperature. Gene expression relative to *Sdha* was calculated with the 2^−ΔCt^ method, with further normalization (as gene expression ratio) on the condition indicated in the figure legend. Primer sequences are indicated in **Table S2**.

### Immunofluorescence analyses

Mouse embryonic samples from timed matings (day of vaginal plug = E0.5) were collected and fixed overnight in 4% paraformaldehyde (PFA), serially dehydrated, embedded in paraffin, and 5-μm-thick transverse sections were prepared. *In vitro*-derived aggregates were processed similarly, with 20 min of fixation and 4% agarose embedding before dehydration, paraffin embedding and sectioning. Cells cultured in 2D were fixed for 10 min in 4% PFA and kept in PBS at 4°C until immunostaining.

Paraffin sections were dewaxed and rehydrated *via* ethanol series. Heat-induced epitope retrieval was performed in sodium citrate buffer pH 6 or Tris-EDTA buffer pH 9. Sections were blocked for 1-2 h at room temperature, and primary antibodies incubated overnight at 4°C. Alexa Fluor-conjugated secondary antibodies were used. Slides were counterstained with Hoechst 33342 (Invitrogen H3570) or DAPI (Thermo Scientific™ 62248) and mounted in PBS:glycerol. Cells on clear-bottom plates were processed similarly with 30 min 0.5% Triton permeabilization instead of heat-induced epitope retrieval. Antibodies and dilutions used are indicated in **Table S3.** Images were acquired using Zeiss LSM780 at the PRISM Facility (Institut de Biologie Valrose, University of Nice Côte d’Azur) and Zeiss ApoTome at CFCI Imaging Facility (TU Dresden) and processed with Zen software (Carl Zeiss) and Fiji (NIH).

### Steroid analysis

The amount of corticosterone released in aggregates’ supernatants was measured by ELISA assay using the Corticosterone rat/mouse ELISA kit (Demeditec #DEV9922) according to the manufacturer’s protocol. Optical density values were determined using a Multiskan™ FC Microplate Photometer (Thermo Scientific™ 51119000).

Furthermore, steroid profiling in cell culture supernatants was performed by LC–MS/MS as previously described ^38^.

### Statistics

Statistical tests were performed using GraphPad Prism 9 (Dotmatics). Unpaired two-tailed Welch’s t-test and one way ANOVA with Welch’s correction were used as indicated. The number of independent experiments performed is indicated (n). All data are presented as the mean ± SEM or ± SD as indicated. p values of <0.05 were considered statistically significant.

## Acknowledgments

We thank Uta Lehnert, Linda Friedrich and Françoise Kühne for technical assistance. The authors greatly acknowledge the teams of the Platform of Resources in Imaging and Scientific Microscopy (PRISM, Institut de Biologie Valrose, University of Nice Côte d’Azur), the Core Facility Cellular Imaging (CFCI, TU Dresden) and the Animal Facility (Faculty of Medicine, University of Geneva), as well as Pauline Sararols for bioinformatic analysis. This work was supported by the Deutsche Forschungsgemeinschaft (DFG, German Research foundation) project no. 314061271, TRR 205/2: “The Adrenal: Central Relay in Health and Disease”, the Fondation pour la Recherche Médicale (grant number ARF201909009270 to Y.N.), La Ligue Contre le Cancer (Equipe Labelisée to A.S.), IFCAH (2022), and the ANR (ANR-11-LABX-0028-01 and ANR-18-CE14-0012 to A.S.).

## Author contributions

Conceptualization: IO, YN & AS ; methodology: IO, YN, MT, FJ, MP ; formal analysis and investigation: IO, YN, MT; data curation: IO, YN; writing – original draft: IO, YN, CS & AS; writing – review & editing: IO, YN, CS & AS; supervision: SN, AS, CS; funding acquisition: YN, SB, SN, CS & AS.

## Declaration of interests

The authors declare no competing interests.

## Notes

### Competing Interest Statement

The authors have declared no competing interest.

## References

1. Walczak, E.M., and Hammer, G.D. (2015). Regulation of the adrenocortical stem cell niche: Implications for disease. Preprint at Nature Publishing Group, 10.1038/nrendo.2014.166 10.1038/nrendo.2014.166.

2. Hellesen, A., Bratland, E., and Husebye, E.S. (2018). Autoimmune Addison’s disease – An update on pathogenesis. Ann Endocrinol (Paris) 79, 157–163. 10.1016/j.ando.2018.03.008.

3. Speiser, P.W., Azziz, R., Baskin, L.S., Ghizzoni, L., Hensle, T.W., Merke, D.P., Meyer-Bahlburg, H.F.L., Miller, W.L., Montori, V.M., Oberfield, S.E., et al. (2010). Congenital Adrenal Hyperplasia Due to Steroid 21-Hydroxylase Deficiency: An Endocrine Society Clinical Practice Guideline. J Clin Endocrinol Metab 95, 4133–4160. 10.1210/jc.2009-2631.

4. White, P.C., New, M.I., and Dupont, B. (1984). HLA-linked congenital adrenal hyperplasia results from a defective gene encoding a cytochrome P-450 specific for steroid 21-hydroxylation. Proc Natl Acad Sci U S A 81, 7505–7509. 10.1073/pnas.81.23.7505.

5. Lightman, S.L., Birnie, M.T., and Conway-Campbell, B.L. (2020). Dynamics of ACTH and Cortisol Secretion and Implications for Disease. Endocr Rev 41, 470–490. 10.1210/endrev/bnaa002.

6. Bornstein, S.R., Malyukov, M., Heller, C., Ziegler, C.G., Ruiz-Babot, G., Schedl, A., Ludwig, B., and Steenblock, C. (2020). New Horizons: Novel Adrenal Regenerative Therapies. J Clin Endocrinol Metab 105, 3103–3107. 10.1210/clinem/dgaa438.

7. Crawford, P.A., Sadovsky, Y., and Milbrandt, J. (1997). Nuclear receptor steroidogenic factor 1 directs embryonic stem cells toward the steroidogenic lineage. Mol Cell Biol 17, 3997–4006. 10.1128/mcb.17.7.3997.

8. Sonoyama, T., Sone, M., Honda, K., Taura, D., Kojima, K., Inuzuka, M., Kanamoto, N., Tamura, N., and Nakao, K. (2012). Differentiation of human embryonic stem cells and human induced pluripotent stem cells into steroid-producing cells. Endocrinology 153, 4336–4345. 10.1210/en.2012-1060.

9. Ruiz-Babot, G., Balyura, M., Hadjidemetriou, I., Ajodha, S.J., Taylor, D.R., Ghataore, L., Taylor, N.F., Schubert, U., Ziegler, C.G., Storr, H.L., et al. (2018). Modeling Congenital Adrenal Hyperplasia and Testing Interventions for Adrenal Insufficiency Using Donor-Specific Reprogrammed Cells. Cell Rep 22, 1236–1249. 10.1016/j.celrep.2018.01.003.

10. Parker, K.L., Rice, D.A., Lala, D.S., Ikeda, Y., Luo, X., Wong, M., Bakke, M., Zhao, L., Frigeri, C., Hanley, N.A., et al. (2002). Steroidogenic factor 1: an essential mediator of endocrine development. Recent Prog Horm Res 57, 19–36. 10.1210/rp.57.1.19.

11. Ikeda, Y., Shen, W.H., Ingraham, H.A., and Parker, K.L. (1994). Developmental expression of mouse steroidogenic factor-1, an essential regulator of the steroid hydroxylases. Mol Endocrinol 8, 654–662. 10.1210/mend.8.5.8058073.

12. Sakata, Y., Cheng, K., Mayama, M., Seita, Y., Detlefsen, A.J., Mesaros, C.A., Penning, T.M., Shishikura, K., Yang, W., Auchus, R.J., et al. (2022). Reconstitution of human adrenocortical specification and steroidogenesis using induced pluripotent stem cells. Dev Cell 57, 2566–2583.e8. 10.1016/j.devcel.2022.10.010.

13. Neirijnck, Y., Sararols, P., Kühne, F., Mayère, C., Weerasinghe Arachchige, L.C., Regard, V., Nef, S., and Schedl, A. (2023). Single-cell transcriptomic profiling redefines the origin and specification of early adrenogonadal progenitors. Cell Rep 42, 112191. 10.1016/j.celrep.2023.112191.

14. Sasaki, K., Oguchi, A., Cheng, K., Murakawa, Y., Okamoto, I., Ohta, H., Yabuta, Y., Iwatani, C., Tsuchiya, H., Yamamoto, T., et al. (2021). The embryonic ontogeny of the gonadal somatic cells in mice and monkeys. Cell Rep 35, 109075. 10.1016/j.celrep.2021.109075.

15. Cheng, K., Seita, Y., Moriwaki, T., Noshiro, K., Sakata, Y., Hwang, Y.S., Torigoe, T., Saitou, M., Tsuchiya, H., Iwatani, C., et al. (2022). The developmental origin and the specification of the adrenal cortex in humans and cynomolgus monkeys. Sci Adv 8, 1–12. 10.1126/sciadv.abn8485.

16. Schnirman, R.E., Kuo, S.J., Kelly, R.C., and Yamaguchi, T.P. (2023). Chapter Five – The role of Wnt signaling in the development of the epiblast and axial progenitors. In Wnt Signaling in Development and Disease, T. P. Yamaguchi and K. B. T.-C. T. in D. B. Willert, eds. (Academic Press), pp. 145–180. 10.1016/bs.ctdb.2023.01.010.

17. Tosolini, M., and Jouneau, A. (2016). From Naive to Primed Pluripotency: In Vitro Conversion of Mouse Embryonic Stem Cells in Epiblast Stem Cells. In Embryonic Stem Cell Protocols, K. Turksen, ed. (Springer New York), pp. 209–216. 10.1007/7651_2015_208.

18. Takasato, M., Er, P.X., Chiu, H.S., Maier, B., Baillie, G.J., Ferguson, C., Parton, R.G., Wolvetang, E.J., Roost, M.S., De Sousa Lopes, S.M.C., et al. (2015). Kidney organoids from human iPS cells contain multiple lineages and model human nephrogenesis. Nature 526, 564–568. 10.1038/nature15695.

19. Lam, A.Q., Freedman, B.S., Morizane, R., Lerou, P.H., Valerius, M.T., and Bonventre, J. V (2014). Rapid and efficient differentiation of human pluripotent stem cells into intermediate mesoderm that forms tubules expressing kidney proximal tubular markers. J Am Soc Nephrol 25, 1211–1225. 10.1681/ASN.2013080831.

20. Mugford, J.W., Sipilä, P., McMahon, J.A., and McMahon, A.P. (2008). Osr1 expression demarcates a multi-potent population of intermediate mesoderm that undergoes progressive restriction to an Osr1-dependent nephron progenitor compartment within the mammalian kidney. Dev Biol 324, 88–98. 10.1016/j.ydbio.2008.09.010.

21. Plachov, D., Chowdhurry, K., Walther, C., Simon, D., Guenet, J.L., and Gruss, P. (1990). Pax8, a murine paired box gene expressed in the developing excretory system and thyroid gland. Development 110, 643–651. 10.1242/dev.110.2.643.

22. Kobayashi, A., Kwan, K.M., Carroll, T.J., McMahon, A.P., Mendelsohn, C.L., and Behringer, R.R. (2005). Distinct and sequential tissue-specific activitites of the LIM-class homeobox gene Lim1 for tubular morphogenesis during kidney development. Development 132, 2809–2823. 10.1242/dev.01858.

23. Sefton, E.M., Gallardo, M., and Kardon, G. (2018). Developmental origin and morphogenesis of the diaphragm, an essential mammalian muscle. Dev Biol 440, 64–73. 10.1016/j.ydbio.2018.04.010.

24. Mahlapuu, M., Ormestad, M., Enerbäck, S., and Carlsson, P. (2001). The forkhead transcription factor Foxf1 is required for differentiation of extra-embryonic and lateral plate mesoderm. Development 128, 155–166. 10.1242/dev.128.2.155.

25. Clipsham, R., Niakan, K., and McCabe, E.R. (2004). Nr0b1 and its network partners are expressed early in murine embryos prior to steroidogenic axis organogenesis. Gene Expression Patterns 4, 3–14. 10.1016/j.modgep.2003.08.004.

26. Häfner, R., Bohnenpoll, T., Rudat, C., Schultheiss, T.M., and Kispert, A. (2015). Fgfr2 is required for the expansion of the early adrenocortical primordium. Mol Cell Endocrinol 413, 168–177. 10.1016/j.mce.2015.06.022.

27. Caneparo, C., Baratange, C., Chabaud, S., and Bolduc, S. (2020). Conditioned medium produced by fibroblasts cultured in low oxygen pressure allows the formation of highly structured capillary-like networks in fibrin gels. Sci Rep 10, 9291. 10.1038/s41598-020-66145-z.

28. Furno, D. Lo, Mannino, G., Pellitteri, R., Zappal, A., Parenti, R., Gili, E., Vancheri, C., and Giuffrida, R. (2018). Conditioned media from glial cells promote a neural-like connexin expression in human adipose-derived mesenchymal stem cells. Front Physiol 9, 1–12. 10.3389/fphys.2018.01742.

29. Drelon, C., Berthon, A., Sahut-Barnola, I., Mathieu, M., Dumontet, T., Rodriguez, S., Batisse-Lignier, M., Tabbal, H., Tauveron, I., Lefrançois-Martinez, A.M., et al. (2016). PKA inhibits WNT signalling in adrenal cortex zonation and prevents malignant tumour development. Nat Commun 7, 12751. 10.1038/ncomms12751.

30. Kulcenty, K., Holysz, M., and Trzeciak, W.H. (2015). SF-1 (NR5A1) expression is stimulated by the PKA pathway and is essential for the PKA-induced activation of LIPE expression in Y-1 cells. Mol Cell Biochem 408, 139–145. 10.1007/s11010-015-2489-9.

31. Kapałczyńska, M., Kolenda, T., Przybyła, W., Zajączkowska, M., Teresiak, A., Filas, V., Ibbs, M., Bliźniak, R., Łuczewski, Ł., and Lamperska, K. (2018). 2D and 3D cell cultures – a comparison of different types of cancer cell cultures. Archives of Medical Science 14, 910–919. 10.5114/aoms.2016.63743.

32. Dedhia, P.H., Sivakumar, H., Rodriguez, M.A., Nairon, K.G., Zent, J.M., Zheng, X., Jones, K., Popova, L. V., Leight, J.L., and Skardal, A. (2023). A 3D adrenocortical carcinoma tumor platform for preclinical modeling of drug response and matrix metalloproteinase activity. Sci Rep 13, 1–12. 10.1038/s41598-023-42659-0.

33. Vidal, V., Sacco, S., Rocha, A.S., Da Silva, F., Panzolini, C., Dumontet, T., Doan, T.M.P., Shan, J., Rak-Raszewska, A., Bird, T. et al. (2016). The adrenal capsule is a signaling center controlling cell renewal and zonation through Rspo3. Genes Dev 30, 1389–1394. 10.1101/gad.277756.116.

34. King, P., Paul, A., and Laufer, E. (2009). Shh signaling regulates adrenocortical development and identifies progenitors of steroidogenic lineages. Proc Natl Acad Sci U S A 106, 21185–21190. 10.1073/pnas.0909471106.

35. Stévant, I., Kühne, F., Greenfield, A., Chaboissier, M.C., Dermitzakis, E.T., and Nef, S. (2019). Dissecting Cell Lineage Specification and Sex Fate Determination in Gonadal Somatic Cells Using Single-Cell Transcriptomics. Cell Rep 26, 3272–3283.e3. 10.1016/j.celrep.2019.02.069.

36. Labrie, F., Luu-The, V., Lin, S.X., Simard, J., and Labrie, C. (2000). Role of 17β-hydroxysteroid dehydrogenases in sex steroid formation in peripheral intracrine tissues. Trends in Endocrinology and Metabolism 11, 421–427. 10.1016/S1043-2760(00)00342-8.

37. Val, P., Martinez-Barbera, J.P., and Swain, A. (2007). Adrenal development is initiated by Cited2 and Wt1 through modulation of Sf-1 dosage. Development 134, 2349–2358. 10.1242/dev.004390.

38. Peitzsch, M., Dekkers, T., Haase, M., Sweep, F.C.G.J., Quack, I., Antoch, G., Siegert, G., Lenders, J.W.M., Deinum, J., Willenberg, H.S., et al. (2015). An LC-MS/MS method for steroid profiling during adrenal venous sampling for investigation of primary aldosteronism. Journal of Steroid Biochemistry and Molecular Biology 145, 75–84. 10.1016/j.jsbmb.2014.10.006.

39. Takasato, M., Er, P.X., Becroft, M., Vanslambrouck, J.M., Stanley, E.G., Elefanty, A.G., and Little, M.H. (2014). Directing human embryonic stem cell differentiation towards a renal lineage generates a self-organizing kidney. Nat Cell Biol 16, 118–126. 10.1038/ncb2894.

40. Shin, E.Y., Park, S., Choi, W.Y., and Lee, D.R. (2021). Rapid Differentiation of Human Embryonic Stem Cells into Testosterone-Producing Leydig Cell-Like Cells In vitro. Tissue Eng Regen Med 18, 651–662. 10.1007/s13770-021-00359-8.

41. Morizane, R., Lam, A.Q., Freedman, B.S., Kishi, S., Valerius, M.T., and Bonventre, J. V. (2015). Nephron organoids derived from human pluripotent stem cells model kidney development and injury. Nat Biotechnol 33, 1193– 1200. 10.1038/nbt.3392.

42. Oeda, S., Hayashi, Y., Chan, T., Takasato, M., Aihara, Y., Okabayashi, K., Ohnuma, K., and Asashima, M. (2013). Induction of intermediate mesoderm by retinoic acid receptor signaling from differentiating mouse embryonic stem cells. International Journal of Developmental Biology 57, 383–389. 10.1387/ijdb.130058ma.

43. Knarston, I.M., Pachernegg, S., Robevska, G., Ghobrial, I., Er, P.X., Georges, E., Takasato, M., Combes, A.N., Jørgensen, A., Little, M.H., et al. (2020). An In Vitro Differentiation Protocol for Human Embryonic Bipotential Gonad and Testis Cell Development. Stem Cell Reports 15, 1–15. 10.1016/j.stemcr.2020.10.009.

44. Cruz Walma, D.A., and Yamada, K.M. (2020). The extracellular matrix in development. Development (Cambridge) 147. 10.1242/dev.175596.

45. Chamoux, E., Bolduc, L., Lehoux, J.G., and Gallo-Payet, N. (2001). Identification of extracellular matrix components and their integrin receptors in the human fetal adrenal gland. Journal of Clinical Endocrinology and Metabolism 86, 2090–2098. 10.1210/jc.86.5.2090.

46. Chamoux, E., Narcy, A., Lehoux, J.G., and Gallo-Payet, N. (2002). Fibronectin, laminin, and collagen IV as modulators of cell behavior during adrenal gland development in the human fetus. Journal of Clinical Endocrinology and Metabolism 87, 1819–1828. 10.1210/jcem.87.4.8359.

47. Otis, M., Campbell, S., Payet, M.D., and Gallo-Payet, N. (2007). Expression of extracellular matrix proteins and integrins in rat adrenal gland: Importance for ACTH-associated functions. Journal of Endocrinology 193, 331–347. 10.1677/JOE-07-0055.

48. Chung, C.Y., Zardi, L., and Erickson, H.P. (1995). Binding of Tenascin-C to Soluble Fibronectin and Matrix Fibrils. Journal of Biological Chemistry 270, 29012–29017. 10.1074/jbc.270.48.29012.

49. Lightner, V.A., and Erickson, H.P. (1990). Binding of hexabrachion (tenascin) to the extracellular matrix and substratum and its effect on cell adhesion. J Cell Sci 95, 263–277. 10.1242/jcs.95.2.263.

50. van Obberghen-Schilling, E., Tucker, R.P., Saupe, F., Gasser, I., Cseh, B., and Orend, G. (2011). Fibronectin and tenascin-C: Accomplices in vascular morphogenesis during development and tumor growth. International Journal of Developmental Biology 55, 511–525. 10.1387/ijdb.103243eo.

51. Ramos, D.M., Chen, B., Regezi, J., Zardi, L., and Pytela, R. (1998). Tenascin-C matrix assembly in oral squamous cell carcinoma. Int J Cancer 75, 680–687. 10.1002/(SICI)1097-0215(19980302)75:5<680::AID-IJC4>3.0.CO;2-V.

52. Chiquet-Ehrismann, R., and Chiquet, M. (2003). Tenascins: Regulation and putative functions during pathological stress. Journal of Pathology 200, 488–499. 10.1002/path.1415.

53. Jones, F.S., and Jones, P.L. (2000). The tenascin family of ECM glycoproteins: Structure, function, and regulation during embryonic development and tissue remodeling. Developmental Dynamics 218, 235–259. 10.1002/(SICI)1097-0177(200006)218:2<235::AID-DVDY2>3.0.CO;2-G.

54. Hu, Y.C., Okumura, L.M., and Page, D.C. (2013). Gata4 Is Required for Formation of the Genital Ridge in Mice. PLoS Genet 9, 1–12. 10.1371/journal.pgen.1003629.

55. Hatano, O., Takakusu, A., Nomura, M., and Morohashi, K.I. (1996). Identical origin of adrenal cortex and gonad revealed by expression profiles of Ad4BP/SF-1. Genes to Cells 1, 663–671. 10.1046/j.1365-2443.1996.00254.x.

56. Heikkilä, M., Peltoketo, H., Leppäluoto, J., Ilves, M., Vuolteenaho, O., Vainio, S., Heikkila, M., Peltoketo, H., Leppäluoto, J., Ilves, M., et al. (2002). Wnt-4 deficiency alters mouse adrenal cortex function, reducing aldosterone production. Endocrinology 143, 4358–4365. 10.1210/en.2002-220275.

57. Butler, A., Hoffman, P., Smibert, P., Papalexi, E., and Satija, R. (2018). Integrating single-cell transcriptomic data across different conditions, technologies, and species. Nat Biotechnol 36, 411–420. 10.1038/nbt.4096.

58. Stallings, N.R., Hanley, N.A., Majdic, G., Zhao, L., Bakke, M., and Parker, K.L. (2002). Development of a transgenic green fluorescent protein lineage marker for steroidogenic factor 1. Molecular Endocrinology 16, 2360–2370. 10.1210/me.2002-0003.

59. Friedrich, L., Schuster, M., Rubin de Celis, M.F., Berger, I., Bornstein, S.R., and Steenblock, C. (2021). Isolation and in vitro cultivation of adrenal cells from mice. STAR Protoc 2, 100999. 10.1016/j.xpro.2021.100999.

60. Nagy, A., Rossant, J., Nagy, R., Abramow-Newerly, W., and Roder, J.C. (1993). Derivation of completely cell culture-derived mice from early-passage embryonic stem cells. Proceedings of the National Academy of Sciences 90, 8424–8428. 10.1073/pnas.90.18.8424.

61. Tamm, C., Galitó, S.P., and Annerén, C. (2013). A comparative study of protocols for mouse embryonic stem cell culturing. PLoS One 8, 1–10. 10.1371/journal.pone.0081156.

