## Supplementary figures for "*In vitro* differentiation of mouse pluripotent stem cells into glucocorticoid-producing adrenocortical cells"

### Supplementary Figure legends:

**Figure S1 (related to Figure 1)**. **Further characterization of the mesodermal differentiation**.

(A) Phase contrast images of cells during the first 8 days of differentiation.

(B-D) RT-qPCR analysis for the EpiSC and PS markers *Fgf5* (*) and *Brachyury* (*T*) (*), respectively (B), the endoderm and paraxial mesoderm markers *Pdx1* (ns) and *Tcf15* (ns), respectively (C), and adrenal and gonadal fated cells *Hoxb9* (****) and *Lhx9* (ns), respectively (D), during the progress of *in vitro* differentiation.

(E) Expression dynamics of *Nr5a1* (**) and *Wt1* (****) until Day10 (total of 6 days in RA+FGF2). Data depict relative mRNA expression, presented as mean ±SEM values of independent replicates (number indicated on the graphs), and normalized on E9.5 mouse urogenital region (E9.5 UR, pools of 4 to 6 embryos).

**Figure S2 (related to Figure 3). Steroidogenic gene expression obtained with different 3D-assembly methods.**

(A-B) RT-qPCR analysis for the steroidogenic genes *Nr5a1, Star, Cyp11a1* and *Cyp21a1* during the progress of *in vitro* differentiation, following aggregation in ULA plates (A) or SP5D (microwell) plates (B). Data depict relative mRNA expression, presented as mean ±SD values of independent replicates (n=3 to 9) and normalized on E14.5 adrenal glands (n=3). Statistical analysis was performed using two-tailed unpaired Welch's t-test. ns.=not significant; *=p < 0.05; **=p < 0.01; ****=p < 0.0001

**Figure S3 (related to Figure 3)**. **Further molecular characterization of aggregates**.

(A-D) RT-qPCR analysis for the zona Glomerulosa gene *Dab2* (A), the gonadal progenitor marker *Lhx9* (B), the testicular steroidogenic marker *Hsd17b3* (C) and the supporting cell markers *Wt1, Sox9, Foxl2* (D) during the progress of *in vitro* differentiation. Data depict relative mRNA expression, presented as mean ±SD values of independent replicates (n=3 to 8) and normalized on adult adrenal cortex+capsule (n=4). Statistical analysis was performed using two-tailed unpaired Welch's t-test. ns, not significant; *=p<0.05; ***=p<0.001.

### Supplementary Tables:

**Table S1**. **Steroid concentrations as measured by LC-MS/MS analysis**

**Table S2. List of primer sequences**

**Table S3. List of antibodies**

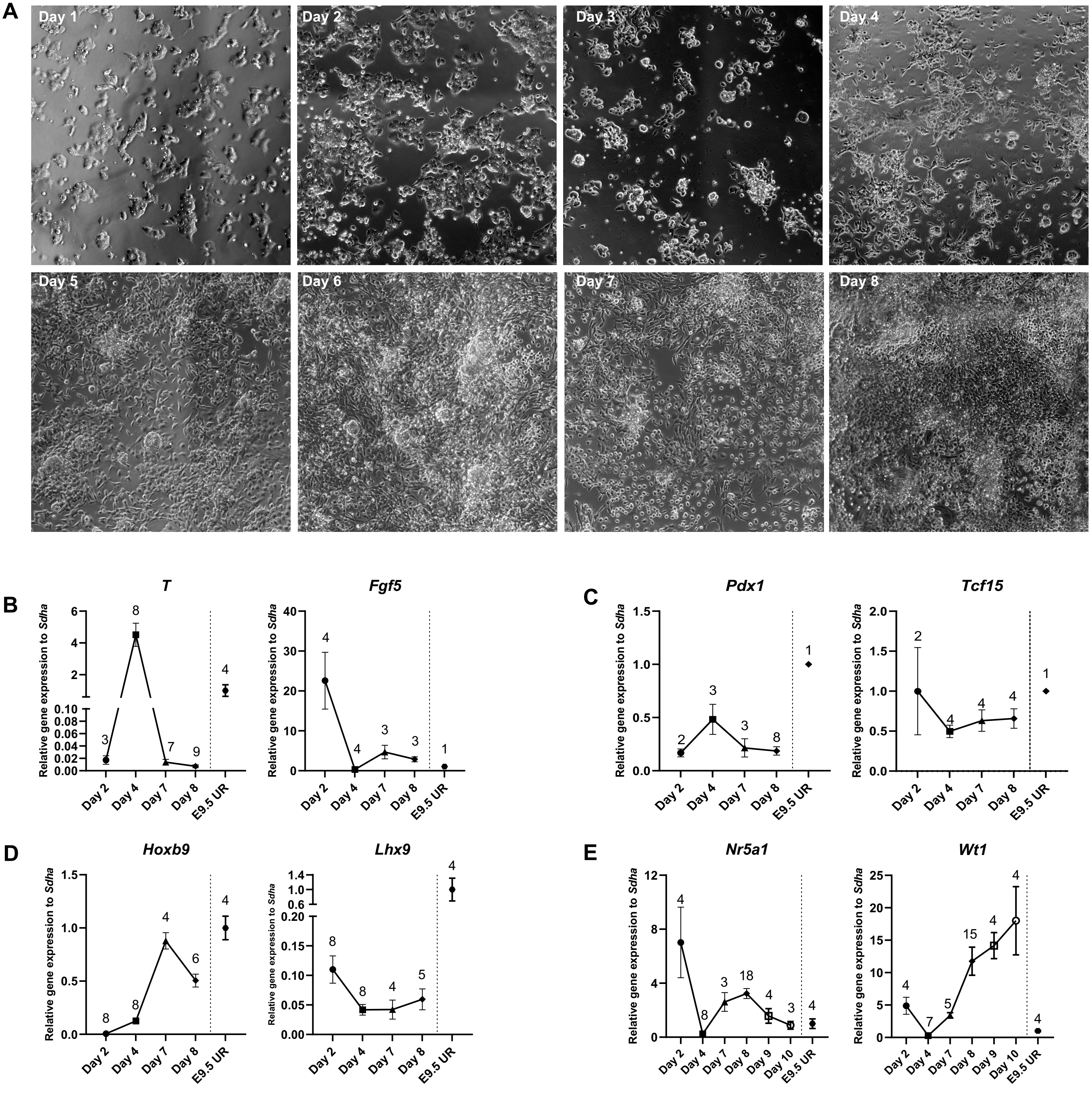


Oikonomakos et al., Fig.S1

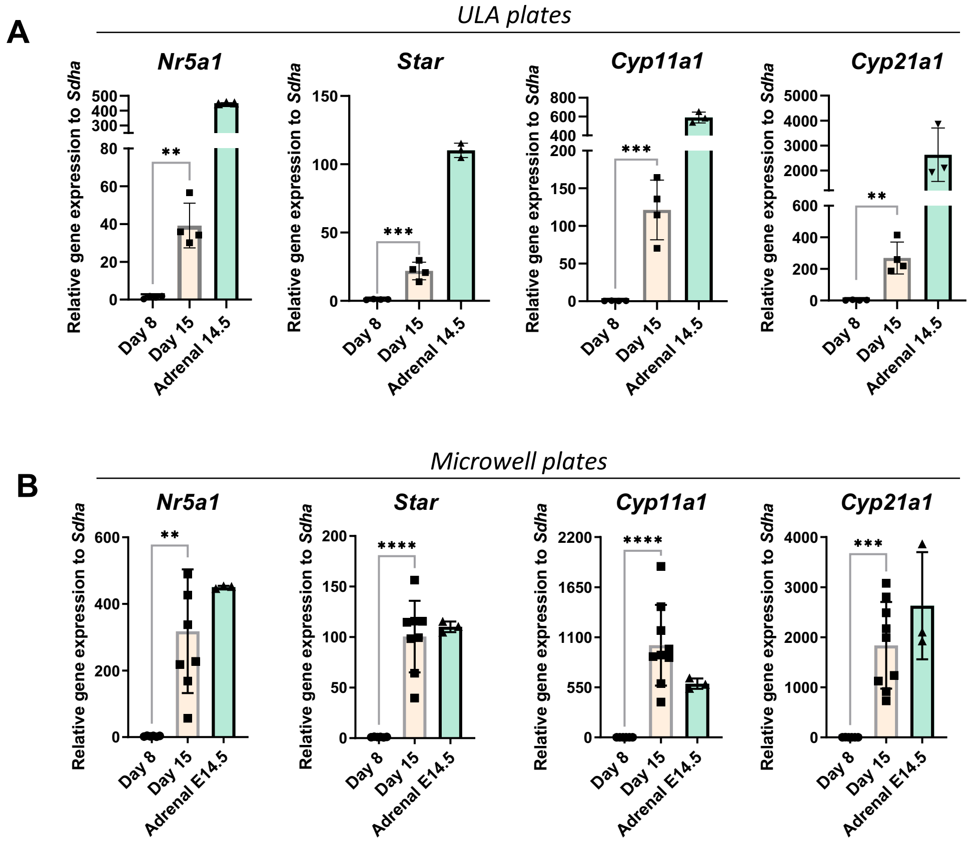


Oikonomakos et al., Fig.S2

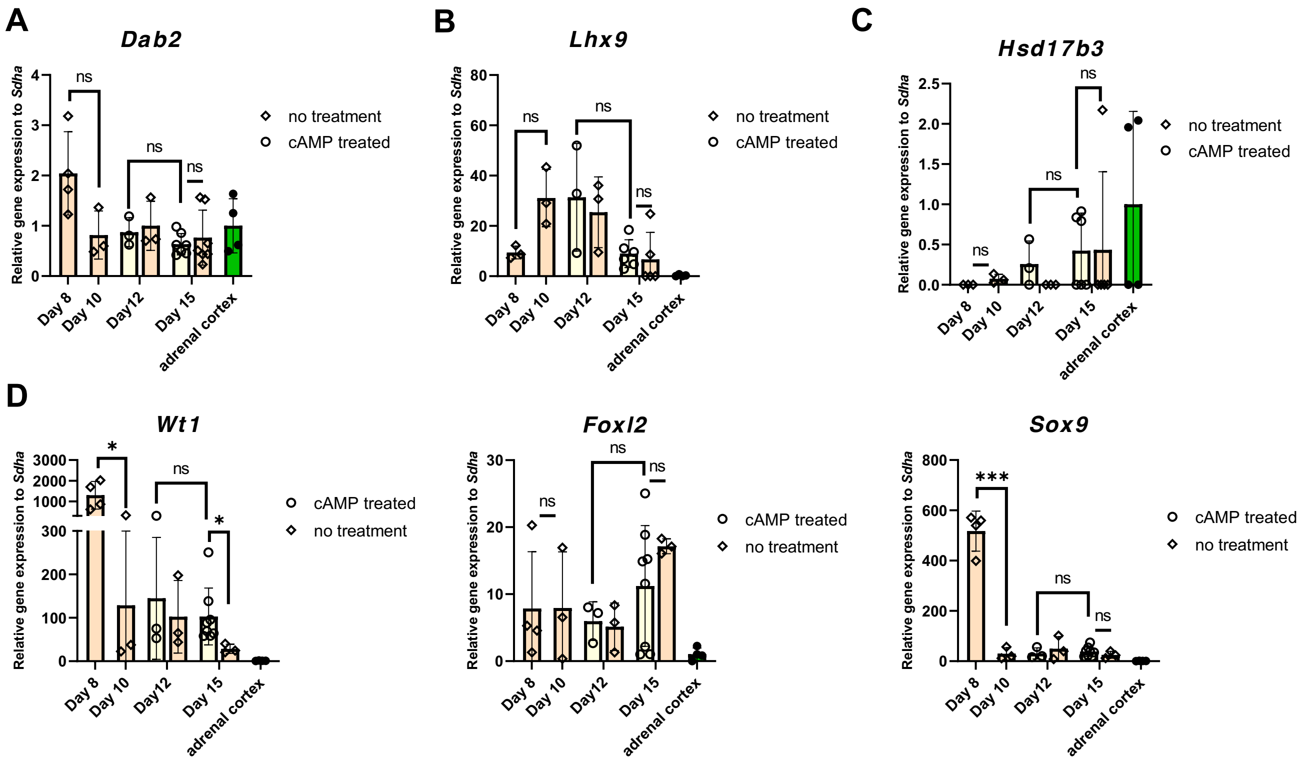


Oikonomakos et al., Fig.S3
